# AIM1-dependent high basal SA accumulation modulates stomatal aperture in rice

**DOI:** 10.1101/2022.06.23.497111

**Authors:** Lei Xu, Hongyu Zhao, Junbin Wang, Xuming Wang, Xianqing Jia, Zhuang Xu, Ruili Li, Kun Jiang, Zhixiang Chen, Jie Luo, Xiaodong Xie, Keke Yi

**Affiliations:** Key Laboratory of Plant Nutrition and Fertilizers, Ministry of Agriculture and Rural Affairs, Institute of Agricultural Resources and Regional Planning, Chinese Academy of Agricultural Sciences, Beijing 100081, China; International Joint Center for the Mechanismic Dissection and Genetic Improvement of Crop Stress Tolerance, College of Agriculture & Resources and Environmental Sciences, Tianjin Agricultural University, Tianjin 300392, China; and College of Basic Sciences, Tianjin Agricultural University, Tianjin 300392, China; State Key Laboratory for Managing Biotic and Chemical Threats to the Quality and Safety of Agro-Products, Ministry of Agriculture Key Laboratory for Plant Protection and Biotechnology, Zhejiang Provincial Key Laboratory of Plant Virology, Zhejiang Academy of Agricultural Sciences, Hangzhou, China; College of Life Sciences, Zhejiang University, Hangzhou, Zhejiang Province, 310058, China; College of Life Sciences, China Jiliang University, 258 Xueyuan Street, Hangzhou, Zhejiang 310018, China, Purdue Center for Plant Biology, Department of Botany and Plant Pathology, Purdue University, 915 W. State Street, West Lafayette, IN 47907-2054, USA; Hainan Key Laboratory for Sustainable Utilization of Tropical Bioresource, Institute of Tropical Agriculture and Forestry, Hainan University, Haikou 570288, China; National Key Laboratory of Crop Genetic Improvement and National Center of Plant Gene Research (Wuhan), Huazhong Agricultural University, Wuhan 430070, China

**Keywords:** Rice, Salicylic acid biosynthesis, OsAIM1, Stomatal, Shoot temperature

## Abstract

The basal levels of salicylic acid (SA), an important plant hormone, vary dramatically among plant species. In the shoot, for example, the monocot plant rice contains almost 100 times higher SA levels than the dicot model plant Arabidopsis. Despite its high basal levels, neither the biosynthetic pathway nor the biological functions of SA is well understood in rice. Here, we report that the synthesis of basal SA in rice shoot is not altered in the mutant for the *ISOCHORISMATE SYNTHASE* (*ICS*) gene, but drastically reduced in the mutant for *OsAIM1*, which encodes a beta-oxidation enzyme in the phenylalanine ammonia-lyase (PAL) pathway. Analogous to its role in thermogenesis, compromised SA accumulation in the *Osaim1* mutant led to a lower shoot temperature than wild-type plants. However, this shoot temperature defect was resulted from increased transpiration due to elevated steady-state stomatal aperture in the mutant. Furthermore, the high basal shoot SA level is required for sustained expression of *WRKY45* to modulate the steady-state stomatal aperture and shoot temperature in rice. Taken together, these results provide the direct genetic evidence for the critical role of the PAL pathway in the biosynthesis of high levels of basal SA, which play an important role in the regulation of steady-state stomatal aperture to promote fitness under both normal and stress conditions.

## Introduction

Salicylic acid (SA) is an important plant hormone that regulates growth, developlent and various stress responses (1). The best established role of SA is the regulation of plant defense response against pathogens infection. The earliest report of SA in plant defense is the effect of exogenously applied SA in protecting tobacco leaves against viral infection (2). Subsequently, it was found that SA could act as an endogenous signal in plant defense (3). During the past 30 years or so, extensive progress has been made in the understanding of the signaling pathway of SA-mediated defense response in plants. In addition to its role in plant defense, SA plays important roles in root growth, leaf senescence and heat production in some thermogenic plants (4, 5).

Endogenous SA is involved in thermogenesis in *Arum lily* flowers for dispersing flower odor to attract pollinators (6). The SA levels in the flowers of the therogenic plants surge about 100-fold upon blooming, leading to a temperature increase of ∼ 10°C above the ambient temperature. Increased levels of SA promotes thermogenesis by inducing the expression of genes encoding alternative oxidase (AOX), a key enzyme in the alternative respiratory pathway in mitochondria that is uncoupled from ATP production and releases energy as heat (7, 8). In most plants, basal SA levels are very low SA (∼ 100 ng/g FW), but they can increase drastically in lily flowers during development or under certain biotic and abiotic stress conditions. Other plants such as populus and rice have very high basal SA levels (>10 μg /g FW) under normal growth conditions. Whether basal SA contributes to thermogenesis in none thermogenic plants, especially those with high basal SA levels, is unknown.

SA is synthesized through two distinct pathways in plants: the isochorismate synthase (ICS) pathway and the phenylalanine ammonia-lyase (PAL) pathway (9). Chorismate is a precursor of both these pathways. ICS catalyzes chorismite to isochorismate in plastid. Isochorismate is exported by ENHANCED DISEASE SUSCEPTIBILITY5 (EDS5) to the cytosol, where the cytosolic amidotransferase avrPphB SUSCEPTIBLE3 (PBS3) catalyzes the conjugation of glutamate to isochorismate to produce isochorismate-9-glutamate, which spontaneously decomposes into SA and 2-hydroxy-acryloyl-*N*-glutamate (Rekhter et al., 2019). On the other hand, PAL catalyzes the chorismate-derived L-phenylalanine into cinnamic acid (CA), which is then converted to SA via o-coumarate or benzoic acid (BA). β-oxidation plays a crucial role in convesion of o-coumarate to SA or cinnamic acid to benzoate (9).

The PAL and ICS biosynthetic routes contribute differently to the biosynthesis of SA in different plant species. *Arabidopsis* contains low basal levels of SA, and most of defense-related SA is synthesized via the ICS pathway (10, 11). A more recent study has showed that ABNORMAL INFLOURESCENCE MERISTEM 1 (AIM1), a peroxinal β-oxiation enzyme, functions in SA biosynthesis in Arabidopsis seeds (12), indicating a role for the AIM1-dependent β-oxidation functions in the PAL pathway in basal SA biosynthesis. In Populus, the single ICS-encoding gene primarily functions in phylloquinone biosynthesis and SA is synthesized from cinnamic acid via the PAL pathway (13, 14). In soybean, the PAL and ICS pathways are equally important for pathogen-induced SA biosynthesis (15). Rice contains a very high basal level of SA, two orders of magnitude higher than *Arabidopsis* (16). However, the contribution of the two biosynthesis pathways to the high basal SA level is not well understood in rice. Early feeding studies suggested that rice shoot converted cinnamic acid and BA into SA (16). In a previous study, we have found that the OsAIM1-dependent PAL pathway is important for SA synthesis in rice roots (17). Rice shoots contain a much higher level of SA tha the roots (18). Although a previous study has implicated ICS in rice SA production (19), there has been no genetic evidence for how SA is produced in rice shoots.

Although rice is well known to contain a higher basal level of SA, it’s biological function is less clear. Unlike in Arabidopsis, SA levels do not greatly increase after pathogen infection in rice (Silverman et al., 1995). However, several studies have suggested that SA plays a role in defense against pathogen infection in rice (20-22). It has been suggested that SA is important for protecting rice from oxidative damage during pathogen infections (23). In addition, we have previously shown that SA is crucial for meristem activity in rice roots. (17) but it is still largely unclear about the biological function of high basal levels of SA in rice shoot.

In this study, we have found that the high basal SA level is unaltered in a rice knockout mutant for its sole ICS gene. By contrast, the mutant for OsAIM1 is drastically reduced in basal SA levels in the rice shoots. These results provide the first genetic evidence for the PAL pathway but against the ICS pathway in the biosynthesis of high basal SA levels in rice. In addition, we have also found that the high basal level of SA in rice shoots is required for maintaining the steady-state stomatal aperture and shoot temperature through a pathway that is dependent on rice WRKY45 transcription factor. We have further shown that the high basal SA levels are important for tolerance to abiotic stresses in rice.

## Results

### OsAIM1 but not OsICS is required for high basal SA accumulation in rice shoot

In plants, SA is synthesized through the ICS pathway and the PAL pathway (Fig. 1*A*). *Arabidopsis* contains low basal levels of SA and syntheize defense-related SA mostly via the ICS pathway. Rice contains much higher levels of SA in shoots but the pathway responsible for rice SA synthesis has not been genetically established. Rice genome contains a single copy *ICS* gene (14). To determine whether the ICS pathway participates in SA biosynthesis in rice shoot, we have isolated and analyzed an *Osics1* mutant (PFG_3A-08161.R) from the T-DNA insertion mutant library (*SI Appendix*, Fig. S1*A*). The mutant was confirmed by PCR-based DNA genotyping using primers flanking the T-DNA insertion site (*SI Appendix*, Fig. S1*B*). The failure to detect the expression of *OsICS1* suggested that *Osics1* is a null mutant (*SI Appendix*, Fig. S1*C*). Importantly, the SA content in the *Osics1* mutant is similar to that in WT (Fig. 1*B*). This result indicates that OsICS1 is not required for high basal levels of SA production in rice shoots.

**Fig. 1.**
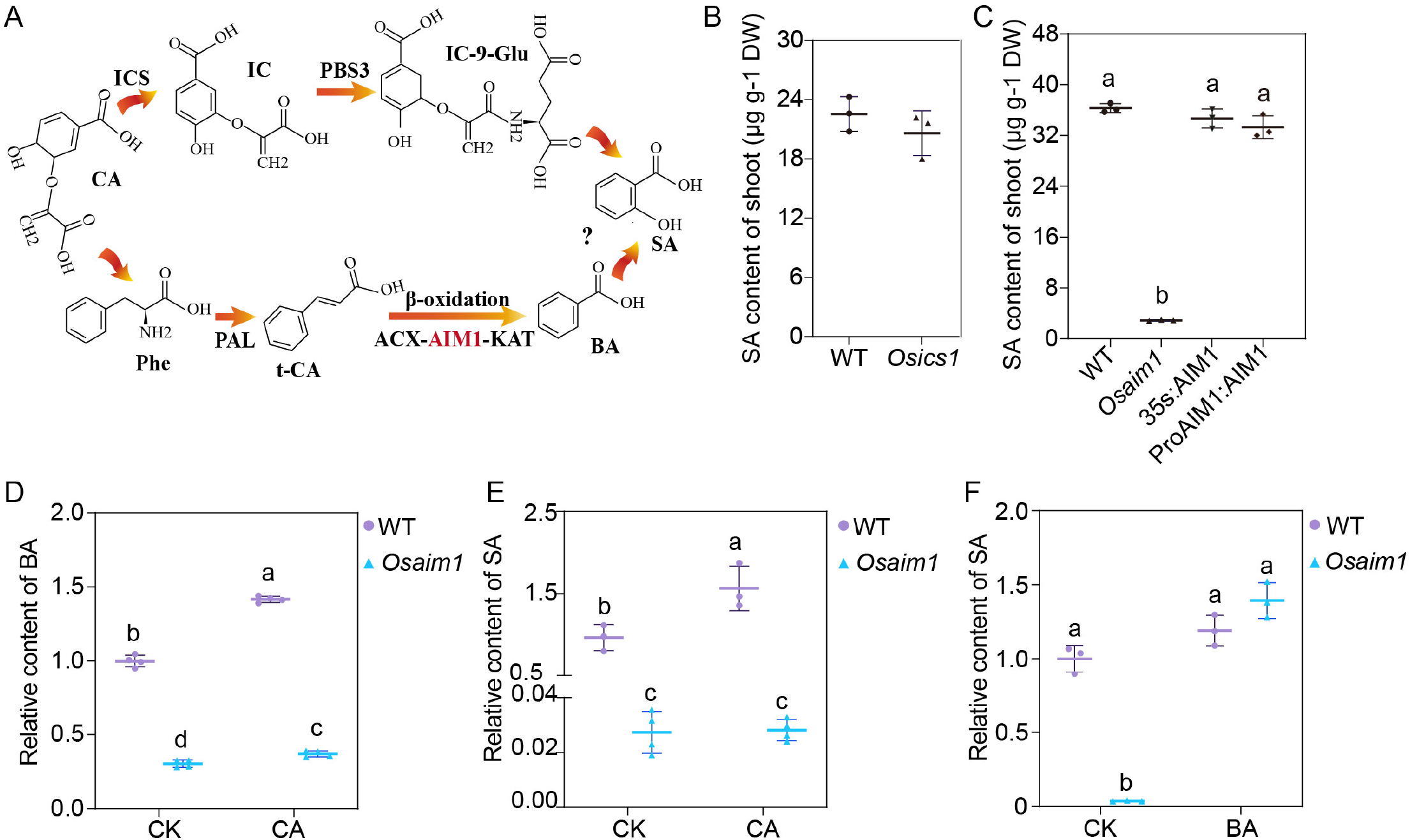
OsAIM1 is essential for SA biosynthesis in rice shoot. **(A)** The biosynthetic pathways for salicylic acid (SA). CA: chorismite, IC: isochorismate, IC-9-Glu, IC-9-glutamate, Phe: Phenylalanine, t-CA: trans-Cinnamic acid,BA: Benzoic acid, SA: Salicylic acid, ICS: isochorismate synthase, PBS3: avrPphB Susceptible 3, PAL: Phenylalanine ammonia lyase, ACX: Acyl-CoA Oxidase, KAT: L-3-Ketoacyl-CoA Thiolase. **(B)** The content of Salicylic acid (SA) in 14-day-old WT and *Osics1* mutants. **(C)** The content of Salicylic acid (SA) in 14-day-old WT, *Osaim1* and complementation lines. 35S indicates complementation lines using 35S promoter, ProAIM1 indicates complementation lines using native promoter. **(D)** The relative content of Benzoic acid (BA) in WT and *Osaim1* plants fed with 50 μM trans-Cinnamic acid (CA) for 7 days. **(E)** The relative content of SA in WT and *Osaim1* plants fed with 50 μM CA for 7 days. **(F)** The relative content of SA in WT and *Osaim1* plants fed with 50 μM BA for 7 days. Error bars represent SD (n = 3 in B, C and F; n=4 in D and E). Different letters show a significant difference by Tukey’s test. The plants were grown for 7 days and then subjected to chemical treatment for 7 days.

In a previous study, we have found that OsAIM1-dependent β-oxidation is required for SA biosynthesis in rice roots and OsAIM1 is also strongly expressed in rice shoots (17). Therefore, OsAIM1 might also participate in SA synthesis in rice shoots. To test this possibility, we measured the shoot SA content in the *Osaim1* mutant. As shown in Fig. 1*C*, the SA content in the mutant shoots was only ∼10% of that in WT shoots. The introduction of the *OsAIM1* gene driven by either its native promoter or the *CaMV 35S* promoter fully restores the SA content of the mutant to the WT levels (Fig. 1*C*). Thus, the OsAIM1-dependent β-oxidation is also required for the high basal levels of SA accumulation in rice shoots.

In the PAL pathway, β-oxidation is required for the shortening of the side chain of CA to produce BA, which is then converted to SA. Consistent with a critical role of OsAIM1 in the β-oxidation of CA in the PAL pathway, the BA levels was also significant reduced in the *Osaim1* mutant whenc compared to those in WT (Fig. 1*D*). To further confirm the role of OsAIM1 in SA biosynthesis, we performed chemical feeding experiments. Application with 50 μM CA resulted in a 1.4-fold increase in both BA and SA levels in WT plants (Fig. 1 *D* and *E*). However, the same feeding treatment in the *Osaim1* mutant only slightly increased the BA levels and had no significant effect on the SA levels (Fig. 1 *D* and *E*). By contrast, feeding with BA restored the SA content of *Osaim1* mutant to the WT levels (Fig. 1*F*). These results demonstrated that OsAIM1-mediated β-oxidation in the PAL pathway is required for the high basal levels of SA production in rice shoots.

### The high basal level of SA is required for modulating the steady-state stomatal aperture and shoot temperature

Rice shoots contain high basal levels of SA with little information on its biological function. In addition to its role in plant defense responses, SA is known to be important for heat production by inducing the expression of alternative oxidases (AOXs) and promoting the alternative respiratory pathway in some thermogenic plants (7). To determine whether the high basal SA content influences the temperature of of rice shoots, we took thermal images of WT and *Osaim1* mutant shoots by infrared thermal camera. Consistently, we observed that the shoot temperature of the *Osaim1* mutant was over 1°C lower than that of WT (Fig. 2 *A* and *B*). The decreased shoot temperature phenotype in *Osaim1* was completely rescued by the *OsAIM1* gene driven by the *CaMV 35S* or *OsAIM1* native promoter (Fig. 2 *A* and *B*). To determine whether reduced SA content is responsible for the observed shoot temperature phenotype in *Osaim1* mutant, we treated WT and *Osaim1* with SA. Exogenous SA application slightly increased the WT shoot temperature and could restore the shoot temperature of the *Osaim1* mutant to the WT level (Fig. 2 *C* and *D*). To determine whether SA elevates rice shoot temperature through upregulating AOX genes as in some thermogeic plants, we compare the *AOX gene* expression between WT and the *Osaim1* mutant. qRT-PCR revealed that AOX gene expression was not compromised but was actually elevated in the *Osaim1* mutant when compared to that in WT (Fig. 2*E*). These findings indicated factors other than the AOX based thermogenesis is responsible for reduced shoot temperature in the *Osaim1* mutant.

**Fig. 2.**
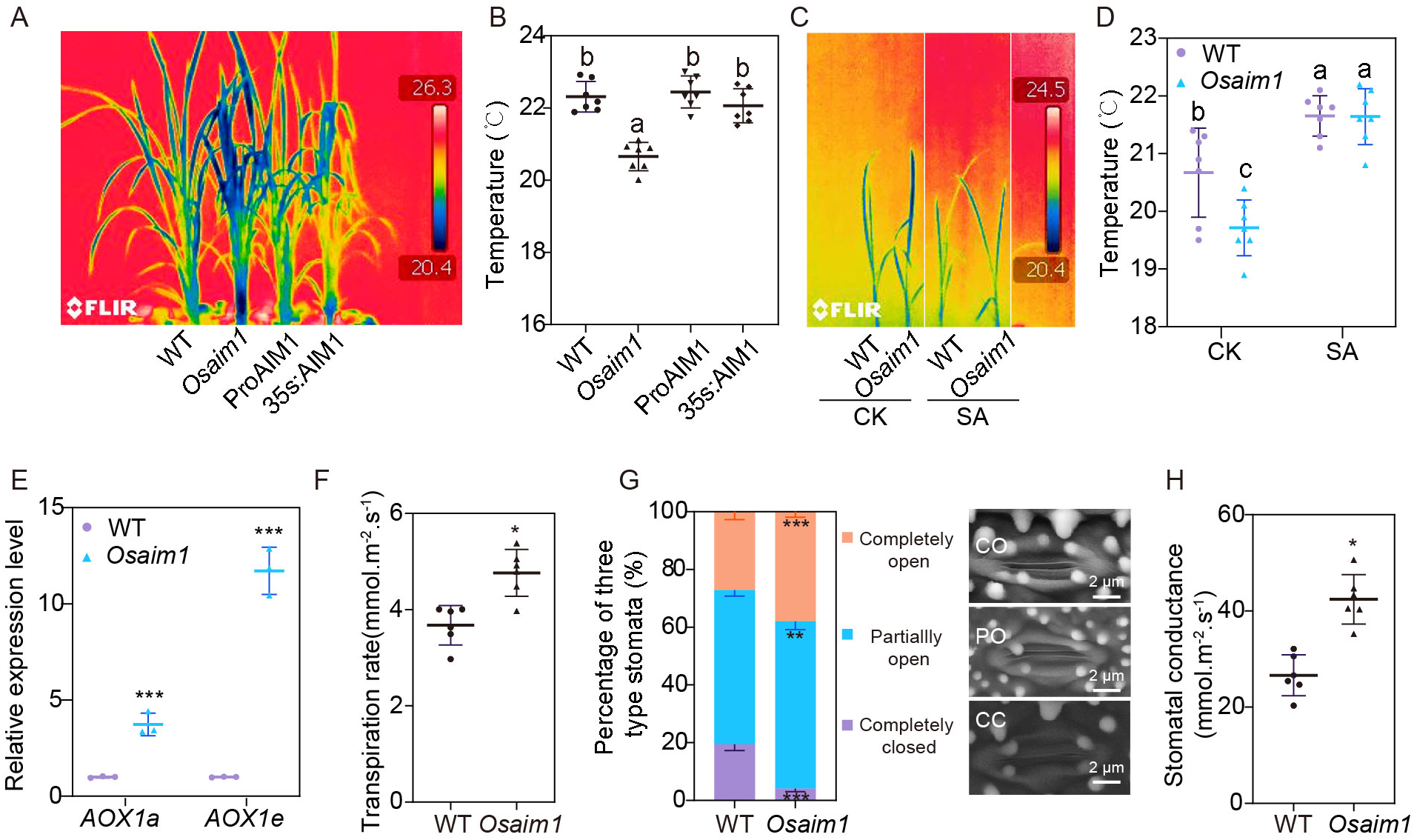
OsAIM1 regulate stomatal aperture and leaf temperature. **(A)** Infrared images of WT, *Osaim1* and complementation lines. 35S indicates complementation lines using 35S promoter, ProAIM1 indicates complementation lines using native promoter. **(B)** Temperature of WT, *Osaim1* and complementation lines. **(C)** Infrared images of WT and *Osaim1* treated with or without SA. **(D)** Temperature of WT and *Osaim1* treated with or without SA. **(E)** The expression level of *AOX1a* and *AOX1e* in WT and *Osaim1*. **(F)** Transpiration rate of WT and *Osaim1*. **(G)**Scanning electron microscopy images of three types of stomatal opening and the percentage of three types of stomatal opening in WT and *Osaim1*. CO: completely open, PO: partially open, CC: completely closed. **(H)** Stomatal conductance of WT and *Osaim1*. Error bars represent SD (n = 7 in B, n = 3 in E, n=6 in D, F and I). Different letters show a significant difference by Tukey’s test. * show a significant difference (* P<0.05, * * P<0.01, ***P<0.001, by Student’s test in E, F and H).

Transpiration also negatively influences leaf temperature (24). Therefore, we examined the potential difference in transpiration between WT and the *Osaim1* mutant. Indeed, the *Osaim1* mutant displayed a transpiration rate approximately 30% higher than WT (Fig. 2*F*). Consistent with increased transpiration, the detached leaves of the *Osaim1* mutant lost water more quickly than WT plants (*SI Appendix*, Fig. S2*A*). Transpiration is mainly determined by stomata density, size and aperture. However, we couldn’t detect any significant difference in the density and size of stomata between WT and *Osaim1* mutant (*SI Appendix*, Fig. S2 *B* and *C*). Therefore, we further checked the stomatal apertures of the WT and *Osaim1* mutant using scanning electron microscop. As shown in Fig. 2*G*, there were higher percentages of completely open stomata in the *Osaim1* mutant (38%) than in WT (27%). The percentage of partially open stomata was about 54% in WT and 58% in the *Osaim1* mutant (Fig. 2*G*). Consistent with the higher percentage of open stomata, the *Osaim1* mutant displayed higher stomatal conductance than WT (Fig. 2*H*).

Taken together, these results strongly suggest that the OsAIM1-dependent PAL pathway is responsible for production of the high basal SA levels in rice shoots, which in turn is required for modulating stomatal aperture and transpiration to influence shoot temperature and other physiological traits.

### SA regulates the stomatal aperture and shoot temperature through a OsWRKY45-dependent pathway

To determine how the high basal SA level regulates stomatal aperture and shoot temperature in rice, we analyzed the factors that are involved in SA signaling. Contrary to the central role of NPR1 in SA signaling in Arabidopsis, SA signaling in rice is mediated by both the OsNPR1 and OsWRKY45 sub-pathways (20, 25). To investigate the roles of OsNPR1 and OsWRKY45 in SA-regulated stomatal aperture and shoot temperature, we generated rice *Oswrky45* and *Osnpr1* loss-of-function mutants by CRISPR-Cas9 (*SI Appendix*, Fig. S3). Like *Osaim1* mutant, the *Oswrky45* mutant displayed significantly lower shoot temperature than WT (Fig. 3 *A* and *B*). Consistent with reduced shoot temperature, the *Oswrky45* mutant had increased transpiration rate than WT plants (Fig. 3*C*). The detached leaves of *Oswrky45* mutant also lost water more quickly than WT plants (*SI Appendix*, Fig. S4*A*). On the other hand, the shoot temperature, transpiration and water loss rate of the *Osnpr1* mutant were all similar to those of WT (Fig. 3 *A*-*C* and *SI Appendix*, Fig. S4*A*). These results suggest that OsWRKY45, but not OsNPR1, mediates SA-regulated stomatal aperture and shoot temperatures. Consistent with this hypothesis, GUS staining of the *P*_*OsWRKY45*_*:GUS* transgenic plant showed that *OsWRKY45* is highly expressed in stomata apparatus (*SI Appendix*, Fig. S5*A*), while expression of *OsNPR1* is absent in stomata apparatus based on the histochemical staining of *P*_*OsNPR1*_*:GUS* transgenic plants (*SI Appendix*, Fig. S5*B*).

**Fig. 3.**
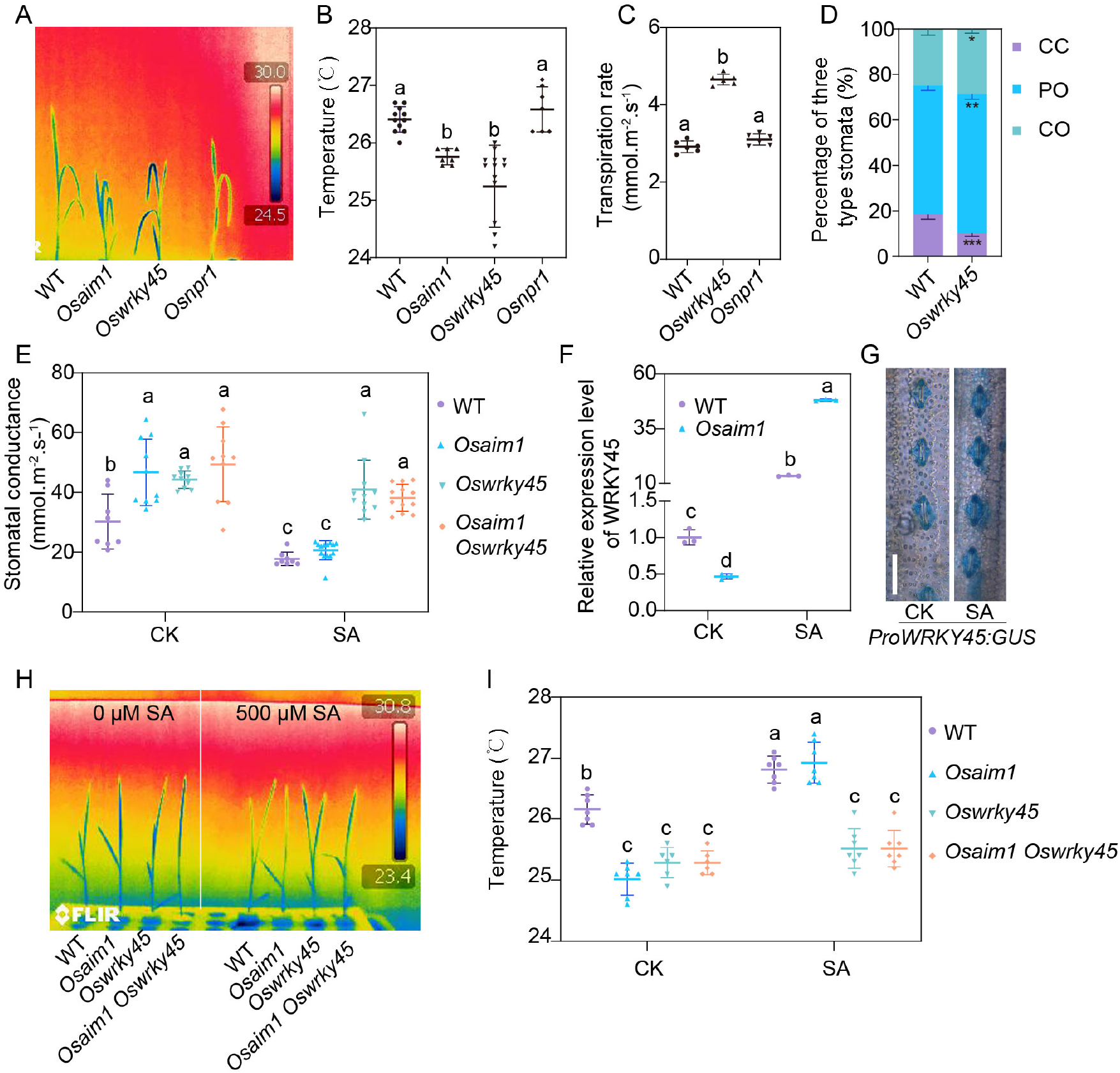
SA derived from OsAIM1 pathway regulate stomatal aperture and leaf temperature through OsWRKY45 dependent pathway. **(A)** Infrared images of WT, *Osaim1, Oswrky45* and *Osnpr1* mutants. **(B)** Temperature of WT, *Osaim1, Oswrky45* and *Osnpr1* mutants. **(C)** Transpiration rate of WT and *Oswrky45*. **(D)** The percentage of three types of stomatal opening in WT and *Oswrky45*. CO: completely open, PO: partially open, CC: completely closed. **(E)**Stomatal conductance of WT, *Osaim1, Oswrky45 and Osaim1 wrky45* treated with or without SA. **(F)** The expression level of *WRKY45* WT and *Osaim1* treated with or without SA. **(G)** The expression of *ProWRKY45:GUS* in stomatal apparatus treated with or without SA treatment. **(H)** Infrared images of WT, *Osaim1, Oswrky45* and *Osaim1 Oswrky45* double mutants treated with 0 or 500 μM SA. **(I)** Temperature of WT, *Osaim1, Oswrky45* and *Osaim1 Oswrky45* double mutants treated with 0 or 500 μM SA. Error bars represent SD (n = 7 in B, n = 5 in C, n=9 in E, n=3 in F, n=7 in I). Different letters show a significant difference by Tukey’s test.

We further analyzed the stomatal apertures of the *Oswrky45* mutant and WT plants using scanning electron microscopy. As shown in Fig. 3*D*, the percentage of completely open stomata in the *Oswrky45* mutant (29%) was significantly higher than that of WT (25%). The percentage of partially open stomata in the *Oswrky45* mutant (61%) was also significantly higher than that of WT (56%) (Fig. 3*D*). As a result, the percentage of completely closed stomata in the *Oswrky45* mutant (10%) was substantially lower than that of WT (19%) (Fig. 3*D*). Consistent with the higher percentage of open stomata, the *wrky45* mutant also displayed higher stomatal conductance than WT (Fig. 3*E*). The decreased shoot temperature and increased stomatal conductance in *Oswrky45* were completely suppressed in the transgenic *Oswrky45* complementation lines containing the *OsWRKY45* genomic fragment (*SI Appendix*, Fig. S4 *B*-*D*). These corroborate the role of *Os*WRKY45 in modulation of stomatal aperture and shoot temperate in rice.

To further determine whether OsWRKY45 acts downstream of SA to regulate stomatal aperture and shoot temperatures, we analyzed the effect of SA on *OsWRKY45* expression. Both the qRT-PCR analysis and GUS staining analysis showed that exogenous SA application strongly induced the expression of *OsWRKY45* in rice shoot and guard cells (Fig. 3 *F* and *G*). To determine the effect of endogenous SA on *OsWRKY45* expression, we further analyzed the expression of *OsWRKY45* in the *Osaim1* mutant and found it to be significantly repressed in the mutant when compared with that in WT (Fig. 3*F*). The exogenous application of SA restored the expression of *OsWRKY45* in the *Osaim1* mutant to the WT level (Fig. 3*F*). By contrast, there was no significant altereation of *OsNPR1* exppression in the *Osaim1* mutant relative to that in WT (*SI Appendix*, Fig. S6). Given that SA treatment could rescue both *OsWRKY45* expression and the defects of stomatal apertures and shoot temperature in the *Osaim1* mutant, it is possible that the high basal SA level in rice shoots promotes *OsWRKY45* expression to regulate stomatal aperture, thereby affecting shoot temperature in rice. Reduced SA levels in the *Osaim1* mutant leads to increased stomatal aperture and reduced shoot temperature due to the reduction of *OsWRKY45* expression.

To test this possibility, we further generated the *Osaim1 Oswrky45* double mutant and compared the effects of SA on the single and double mutants. First, the *Osaim1 Oswrky45* double mutant had a shoot temperature similarily lower than WT as the *Osaim1* or *Oswrky45* single mutant (Fig. 3 *H* and *I*). Thus, the effects of the *Osaim1* and *Oswrky45* mutations are not additive, supporting that they act in the same pathway. Secondly, unlike the *Osaim1* mutant, whose reduced shoot temperature could be restored by SA treatment, the *Oswrky45* single and *Osaim1 Oswrky45* double mutants were insensitive to SA application for restoration of their reduced shoot temperature (Fig. 3 *H* and *I*). Likewise, while SA fully suppressed the elevated stomatal conductance of the *Osaim1* mutant, it failed to do so with the *Oswrky45* single and *Osaim1 Oswrky45* double mutants (Fig. 3*J*). The non-additive nature of the effects of the *Osaim1* and *Oswrky45* mutations and their differential responses to SA strongly support that OsWRKY45 acts downstream of the high basal shoot SA levels in the regulation of stomatal apertures and shoot temperatures in rice.

### The high rice shoot basal SA maintains OsWRKY45 expression to affect H_2_O_2_ accumulation in stomatal apparatus and promote abiotic stress tolerance

H_2_O_2_ is an important signaling molecule involved in the regulation of stomatal aperture (26). Given the critical role of OsAIM1 and OsWRKY45 in the regulation of stomatal apertures, we further investigated the H_2_O_2_ levels in the *Osaim1* and *Oswrky45* mutants using 3^/^-(ρ-hydroxyphenyl) fluorescein (HPF) staining. Indeed, the H_2_O_2_ levels in the guard cells of the *Osaim1, Oswrky45* and *Osaim1 Oswrky45* double mutants were decreased when compared to those in WT. To determine whether H_2_O_2_ is involved in OsWRKY45-regulated stomatal aperture, we also determined the effect of SA on the H_2_O_2_ levels in the guard cells of WT, *Osaim1, Oswrky45* and *Osaim1 Oswrky45*. SA treatment stimulated H_2_O_2_ production in WT and could rescue the defect of reduced H_2_O_2_ accumulation in the *Osaim1* mutant, but not in the *Oswrky45* single and *Osaim1 Oswrky45* double mutants (Fig. 4 *A* and *B*). These results indicate that the high basal SA level enhances H_2_O_2_ accumulation to regulate stomatal aperture in a OsWRKY45-dependent manner.

**Fig. 4.**
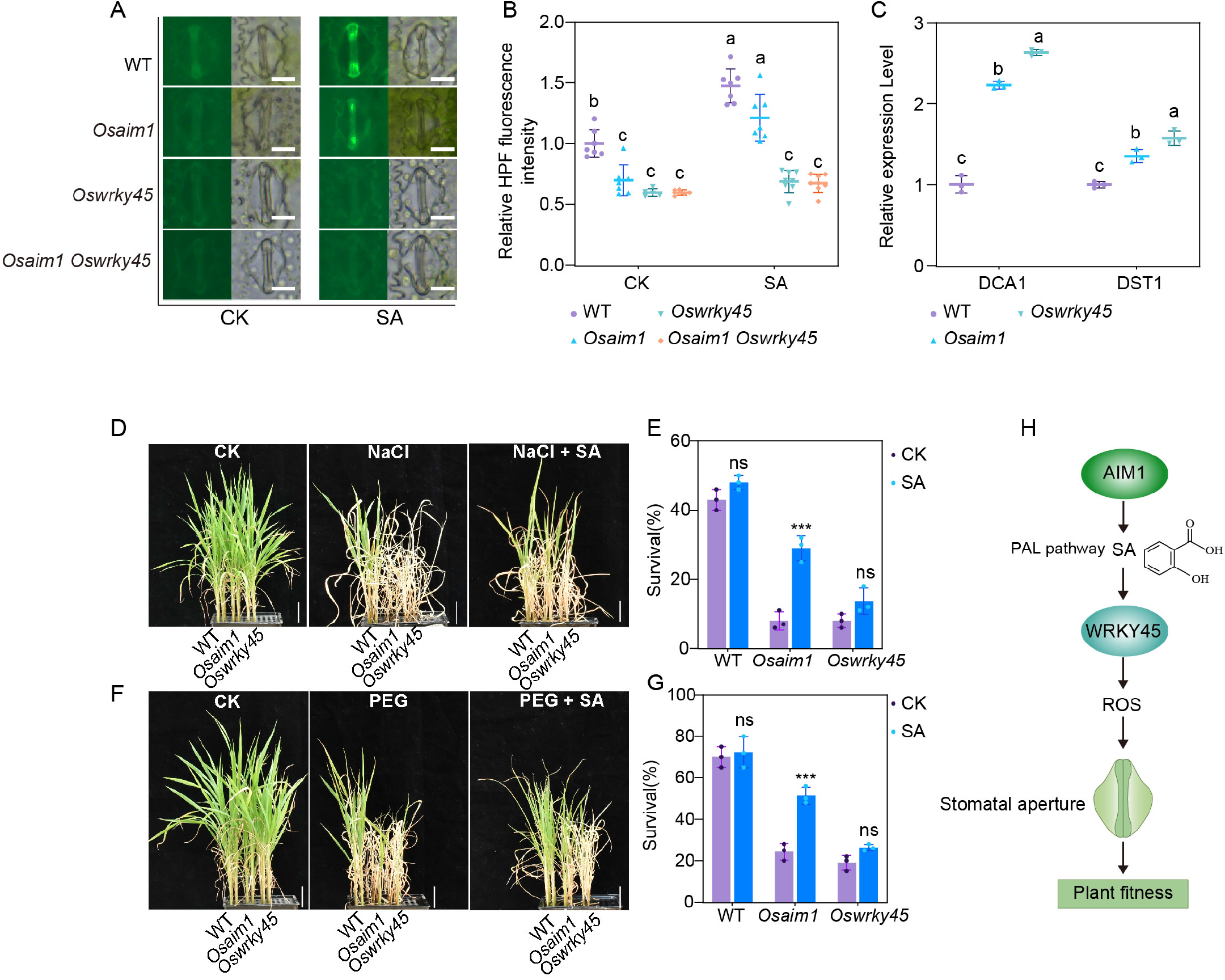
SA regulates ROS accumulation, stomatal aperture and plant fitness in a OsWRKY45 dependent manner. **(A)** Images of H_2_O_2_-based fluorescence signal in guard cells of 21-day-old seedlings of the WT, *Osaim1, Oswrky45* and *Osaim1 Oswrky45* double mutants. **(B)** Intensity of H_2_O_2_-based fluorescence signal in guard cells. The fluorescence intensity was quantified as the average pixel intensity of each guard cell using Image J. **(C)** The expression level of *DCA1* and *DST* in WT, *Osaim1* and *Oswrky45*. **(D)** Salt stress treatment of WT, *Osaim1* and *Oswrky45* mutant. The 14-day-old seedlings were treated with 100 mM NaCl and 100 mM NaCl + 200 µM SA for 7d (SA was added before NaCl treatment for 6h) and then recovered for 10 d. **(E)** The survival rate of WT, *Osaim1* and *Oswrky45* plants recovered for 7 d after NaCl treatment with or without SA treatment. **(F)** PEG 6000 treatment of WT, *Osaim1* and *Oswrky45* mutant. The 14-day-old seedlings were treated with 20% PEG and 20% PEG + 200 µM SA for 10d (SA was added before PEG treatment for 6h) and then recovered for 10 d. **(G)** The survival rate of WT, *Osaim1* and *Oswrky45* plants recovered for 7 d after PEG treatment with or without SA treatment. **(H)** A working model for the role of the OsWRKY45 mediated SA signaling in rice. SA was produced through OsAIM1 dependent PAL pathway in rice shoot. WRKY45-ROS acts downstream of SA to regulate stomatal aperture and abiotic stress response. Error bars represent SD (n = 7 in B, n = 3 in C, n=9 in E, n=3 in E and G). Different letters show a significant difference by Tukey’s test. * show a significant difference (***P<0.001, by Student’s test).

In rice, the DST zinc finger protein transcription factor interacts with another zinc finger protein DCA1 as co-activator to affect drought and salt tolerance in rice via stomatal aperture control by regulating H_2_O_2_ homeostasis in rice (27, 28). We analyzed the expression of *OsDST* and *OsDCA1* in both the *Osaim1* and *Oswrky45* mutants and found expression levels of both genes to be increased in the *Osaim1* and *Oswrky45* mutants (Fig. 4*C*). These results suggest that DCA1 and DST may act downstream of OsWRKY45 to affect H_2_O_2_ accumulation and stomatal aperture in rice.

Steady-state stomatal aperture is crucial for plant adaptation to abiotic stress (27, 29, 30) and, therefore, the positive regulation of OsWRKY45 expression by the high basal shoot SA content could play a role in the plant tolerance to abiotic stress in rice. To test this, we first assessed the responses of WT and the *Osaim1* to salt and polyethylene glycol (PEG) treatments. Fourteen-day-old seedlings were first treated with 100 mM NaCl for 7 days or 20% PEG 6000 for 10 days and then recovered in the normal growth medium for 10 days. As shown in Figure 4D, the *Osaim1* mutant was substantially more sensitive to the salt and PEG treatments than WT. Approximately 40% and 70% of WT plants survived after the salt and PEG treatments, respectively (Fig. 4 *D*-*G*). For comparison, only 8% and 24% of the *Osaim1* mutant plants survived under the salt and PEG treatments, respectively (Fig. 4 *D*-*G*). Furthermore, exogenous application of SA could significantly increase the survival rate of *Osaim1* under the salt and PEG treatments. Thus, compromised SA synthesis resulted in hypersensitivity to salt and PEG treatments in the *Osaim1* mutant. Similar to the *Osaim1* mutant, the *Oswrky45* mutant was less tolerant to both the salt and PEG treatments. However, SA application failed to restore the tolerance to salt and PEG treatments in the *Oswrky45* mutant (Fig. 4 *D*-*G*).

Taken all these results together, OsAIM1-dependent PAL pathway is essential for the accumulation of high basal level SA in rice shoot. In addition, the high basal level of SA is required for the regulation of the steady-state stomatal aperture through a OsWRKY45-dependent pathway in rice, which is crucial for abiotic stresses tolerance in the important crop (Fig. 4*H*).

## Discussion

SA acts as a signaling molecule in plant inducible responses against various forms of environmental stresses and its biosynthesis is stress-responsive in many plant species. SA can be synthesis through two distinct pathways in plants, the ICS pathway and the PAL pathway (31). ICS plays an important role in SA production in some plant species (32). In *Arabidopsis thaliana, AtICS1* is essential for stress-induced SA production. Likewise, a *Nicotiana benthamiana* ICS gene (*NbICS*) is transcriptionally induced by a pathogen elicitor and is required for induced SA production in response to biotic and abiotic stress conditions (33, 34). In contrast, ICS in tobacco (*Nicotiana tabacum*; *NtICS*) does not appear to be involved in stress-induced SA production. The transcript levels of *NtICS* were not induced, and ICS activity was undetectable when SA production is induced by tobacco mosaic virus inoculation or ozone exposure (35). Rice constitutively contains two to three orders of magnitude higher levels of SA than *Arabidopsis, N. benthamiana*, and tobacco. If OsICS is required for production of such higher levels of SA in rice, its activity would be expected to be much higher than that of AtICS1, NbICS, and NtICS. However, a recent study has shown that the *in vitro* and *in planta* ICS activities of OsICS were much lower than those of AtICS1 (36). Previously, we have found that OsAIM1-dependent β-oxidation is required for SA biosynthesis in rice roots (17). In this study, we have further found that OsAIM1 is also required for SA production in the shoots. The basal SA level in the shoots of rice *Osaim1* mutant was reduced by almost 90% when compared to that in WT shoots. Thus, OsAIM1-dependent β-oxidation in the PAL pathway is responsible for synthesis of a majority of basal SA in rice.

In Arabidopsis, most of pathogen-induced SA is derived from isochorismate, which is generated from chorismate by ICS1 in the plastid (32). Isochorismate generated by ICS is also an early precursor for phylloquinone, which can be synthesized via 1,4-dihydroxy-2-naphthoic acid (DHNA) in a multistep pathway (37). As a result, Arabidopsis *ics1 ics2* double mutant is deficient not only in SA but also in phylloquinone biosynthesis. Because phylloquinone is essential for electron transfer in photosystem I, the *ics1 ics2* double mutants are smaller than WT and display chlorosis (11). Like rice, barley contains a single ICS gene. Phylloquinone in the barley *ics1* mutant is undetectable and the mutant displays wilting symptoms, both of these phenotypes could be rescued by DHNA application, indicating that the wilting phenotype of barley *ics1* mutant is caused by phylloquinone deficiency (38). However, the SA level in barley *ics1* mutant was comparable to that in WT. Similarly, we have found that the SA content in the *Osics1* mutant was. similar to that of WT. These results indicate that OsICS1 is required for production of phylloquinone but not SA in rice.

Rice shoot accumulates an extremely high basal level of SA, almost 100-fold higher than that in Arabidopsis. A previous report has found that SA plays an important role in protecting rice from oxidative damage (23). Here, we found that the high basal SA level in rice shoot is required for the maintenance of shoot temperatures by modulating stomatal aperture. The *Osaim1* mutant had about 90% less SA than WT in shoot and displayed increased stomatal aperture and reduced shoot temperatures relative to those in WT. NPR1 is the master regulator of SA-mediated responses (39) and SA induces stomatal closure in an NPR1-dependent manner in Arabidopsis (40). In rice, however, both OsNPR1 and OsWRKY45 play a crucial role in SA signaling (20). Our results demonstrate that OsWRKY45, rather than OsNPR1, plays a critical role in SA-regulated stomatal aperture. H_2_O_2_ is an important signal molecule that induces stomatal closure (26). We have found that OsWRKY45-dependent SA signaling pathway may control stomata aperture by regulating H_2_O_2_ accumulation. H_2_O_2_ level was decreased in the *Osaim1, Oswrky45* single and *Osaim1 Oswrky45* double mutants. SA treatment could rescue the H_2_O_2_ accumulation defect of *Osaim1*, but not the *Oswrky45* and *Osaim1 Oswrky45* mutants. Furthermore, we found that the expression of *OsDCA1* and *OsDST*, negative regulators of stomatal closure via regulating H_2_O_2_ homeostasis, increased in the *Osaim1* and *Oswrky45* mutants. These results suggest that high basal SA level in rice shoot may maintain the steady-state stomatal aperture through OsWRKY45-OsDST/OsDCA1-H_2_O_2_ signaling pathway.

In addition to its crucial role in plant biotic stress response, SA plays an important role in plant response to abiotic stresses, such as salt, drought and cold (1). Although many studies have analyzed the effect of SA on abiotic stresses in plants, its role and molecular mechanism in abiotic stress tolerance in rice is still poorly understood. Steady-state stomatal aperture is a key component in plant adaption to abiotic stress (27, 29, 30). Therefore, the high basal SA level in rice shoot may maintain the steady-state stomatal aperture to increase the abiotic stress tolerance. Indeed, we have found that *Osaim1* and *Oswrky45* are more sensitive to NaCl and PEG treatment, and exogenous SA could restore this phenotype of *Osaim1*, but not *Oswrky45*. These results support that steady-state stomatal aperture maintained by the OsWRKY45-dependent SA signaling pathway is crucial for abiotic stress tolerance in rice. As sessile organisms constantly exposed to diverse environmental challenges, including biotic and abiotic stresses, plants have evolved complex molecular mechanisms to increase their adaptation and fitness. In this study, we have established that OsWRKY45 is a key player in SA-mediated abiotic stresses adaptation. Previous studies have found that WRKY45 plays a crucial role in SA-mediated defense signaling by activating redox-related genes in rice (20). Regulation of stomatal aperture also contributes to plant defense by restricting pathogen entry into plants (41). Thus, OsWRKY45 is a critical regulator of stomatal closure in the SA signaling pathway in plant responses to both biotic and abiotic stresses.

In summary, we have showed here that the OsAIM1-dependent PAL pathway, not the ICS pathway, is essential for biosynthesis of high basal levels of SA in rice shoot. We have also found that the high basal SA level in rice shoot is required for maintaining the steady state stomatal aperture through a OsWRKY45 dependent pathway, which is important for enhanced plant tolerance to abiotic stresses.

## Material and Methods

### Plant Materials and Growth Conditions

Hydroponic experiments were conducted using a modified rice culture solution containing 1.425 mM NH_4_NO_3_, 0.2 mM NaH_2_PO_4_, 0.513 mM K_2_SO_4_, 0.998 mM CaCl_2_, 1.643 mM MgSO_4_, 0.009 mM MnCl_2_, 0.075 mM (NH_4_)_6_Mo_7_O_24_, 0.019 mM H_3_BO_3_, 0.155 mM CuSO_4_, and 0.152 mM ZnSO_4_ with 0.125 mM EDTA-Fe. pH of the solution was adjusted to 5.5. All the plants were grown in a greenhouse with a 12 h day (30 °C)/12 h night (22 °C) photoperiod, approximately 200 μmol m^-2^ s^-1^ photon density, and approximately 60 % humidity.

### Plasmid Construction and Plant Transformation

The sequence information for primers used to construct vectors is shown in Supplemental Table 1 online. For *ProWRKY45:GUS* vector, the 2753 bp promoter before start code was introduced into the PstI and KpnI sites on pCAMBIA 1300-GUS vector. For *ProNPR1:GUS* vector, the 2425 bp promoter before start code was introduced into the SalI and KpnI sites on pCAMBIA 1300-GUS vector. For *ProWRKY45:WRKY45* vector, the 4428 bp genome fragment was introduced into the KpnI and XbaI sites on pCAMBIA 1300 vector. The constructs were transformed into mature embryos developed from seeds of WT (‘Shishoubaimao’) or *wrky45* mutant via *Agrobacterium tumefaciens* mediated transformation.

### Histochemical Localization of GUS Expression

For GUS staining, the tissues were incubated in a solution containing 50 mM sodium phosphate buffer (pH 7.0), 5mM K3Fe(CN)6, 5 mM K4Fe(CN)6, 0.1% Triton X-100, and 1 mM X-Gluc at 37?. Sections (35μm) of various plant tissues were made by vibratome (Leica VT 1000 S). Images were taken by microscope (Nikon ECLIPSE Ni).

### Quantitative RT-PCR

Reverse transcription was performed using 2 µg of total RNA and M-MuLV Reverse Transcriptase (NEB) according to the manufacturer’s instructions. Quantitative PCR (qPCR) was performed using the Roche SYBR green I kit on the LightCycler480 machine (Roche Diagnostics) according to the manufacturer’s instructions. Three biological replicates were performed for each gene. Rice *Actin* gene was used as an internal control.

### NBT and HPF staining

Rice Roots were stained for 30 min in a solution of 2 mM nitroblue tetrazolium (NBT) or 5 µM 3’
s-(p -hydroxyphenyl) fluorescein (HPF) in 20 mM phosphate buffer (pH 6.1). The reaction was stopped by transferring the seedlings in distilled water. Root tips were imaged under bright-field illumination for NBT staining or GFP channel for HPF staining on a Nikon microscope. The Intensity of NBT and HPF staining was quantified using ImageJ software.

### Metabolites measurement

For metabolites measurements, seedlings were grown in cultural solution for two weeks. The aerial parts and roots were separately collected and freeze-dried. The freeze-dried tissues were used for Metabolites measurement, following previously published methods (Chen et al., 2013). The sample extracts were analyzed using an LC–ESI– MS/MS system (HPLC, Shim-pack UFLC SHIMADZU CBM20A system, www.shimadzu.com; MS, Applied Biosystems 4000 Q TRAP, www.appliedbiosystems.com).

### Stomatal Aperture Measurement

For imaging rice stomata, leaves of 3-week-old plants were fixed with 2.5% (v/v) glutaraldehyde, and stomatal pictures were obtained by a Hitachi TM3030Plus scanning electron microscopy. At least 60 stomata were examined in each line, and the assays were repeated three times.

### Measurements of water loss

Water loss rates of the detached leaves from 3-week-old plants were measured by monitoring the fresh weight loss at indicated points under a constant temperature (28 °C) and humidity (60 %). The percentage loss of weight was calculated based on the initial weight of the plants.

### Thermal Imaging

Thermal images of the plant were taken with an infrared thermal camera (FLIR T400; FLIR Systems, Boston, MA, USA) and were analyzed subsequently through the FLIR R&D software.

### Gas Exchange Measurements

The stomatal conductance (Gs) and transpiration rate (E) were measured using a LI-6800 portable photosynthesis system (PP Systems, USA) at 28 °C, 200 μmol m^−2^ s^−1^ light, 400 μmol mol^−1^ CO_2_ and 60 % relative humidity.

## Acknowledgements

This work was supported by the Central Public-interest Scientific Institution Basal Research Fund (No. Y2022QC14), the National Natural Science Foundation (32130096, 31800246) and Natural Science Foundation of Tianjin (20JCYBJC00260). K.Y. was supported by the Agricultural Science and Technology Innovation Program of Chinese Academy of Agricultural Sciences.

## Competing interests

The authors declare no competing interests.

## Notes

### Competing Interest Statement

The authors have declared no competing interest.

