## Supplemental Figure1-6 and Supplemental Table 1 for "AIM1-dependent high basal SA accumulation modulates stomatal aperture in rice"

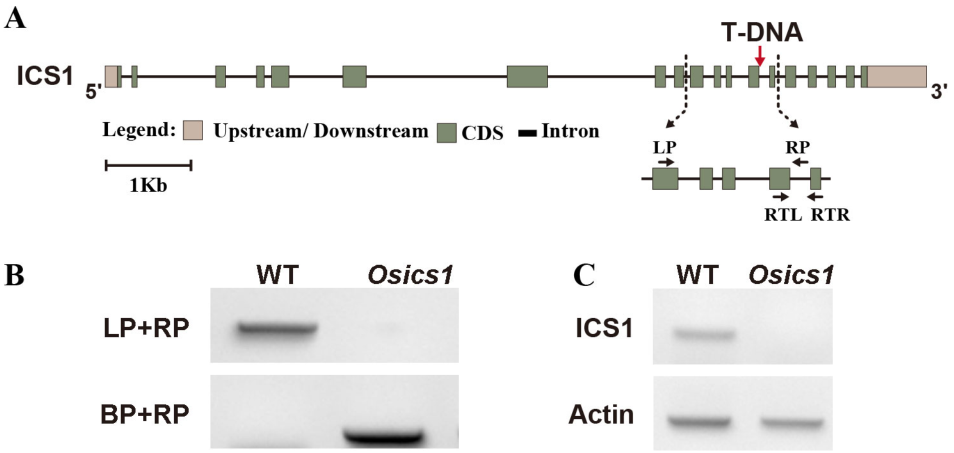

**Fig. S1 Characterization of *Osics1* mutant.**

**(A)** Schematic representation of *OsICS1*. White boxes indicate the untranslated Regions, black boxes indicate the CDS and lines represent the introns. The black arrows indicate the insertion sites of T-DNA. **(B)** Verification of the T-DNA insertion by PCR in the *Osics1* mutant. The positions of the F, R, and BP primers are indicated as red arrows. **(C)** Verification of the T-DNA insertion by RT-PCR in the *Osics1* mutant. The positions of the RTF and RTR primers are indicated as red arrows.

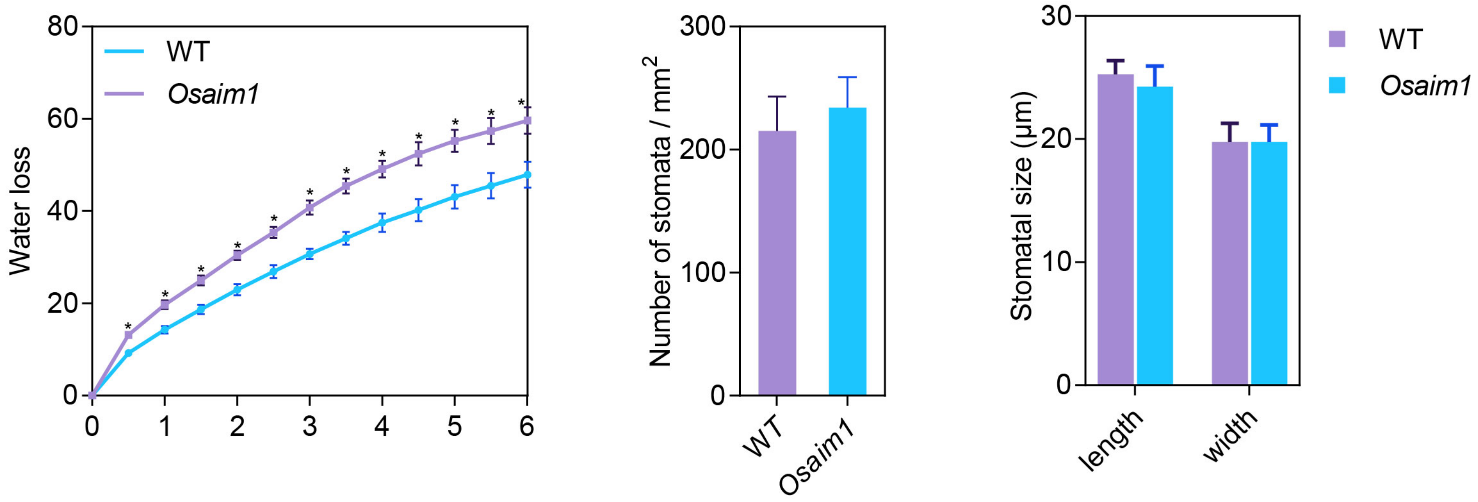

**Fig. S2** Water loss, stomata density and size of WT and *Osaim1*.

**(A)** Water loss of detached leaves of WT and *Osaim1*. **(B)** Stomatal density of WT and *Osaim1*. **(C)** Stomatal size of WT and *Osaim1*. Error bars represent SD. * show a significant difference (*P<0.05, by Student's test).

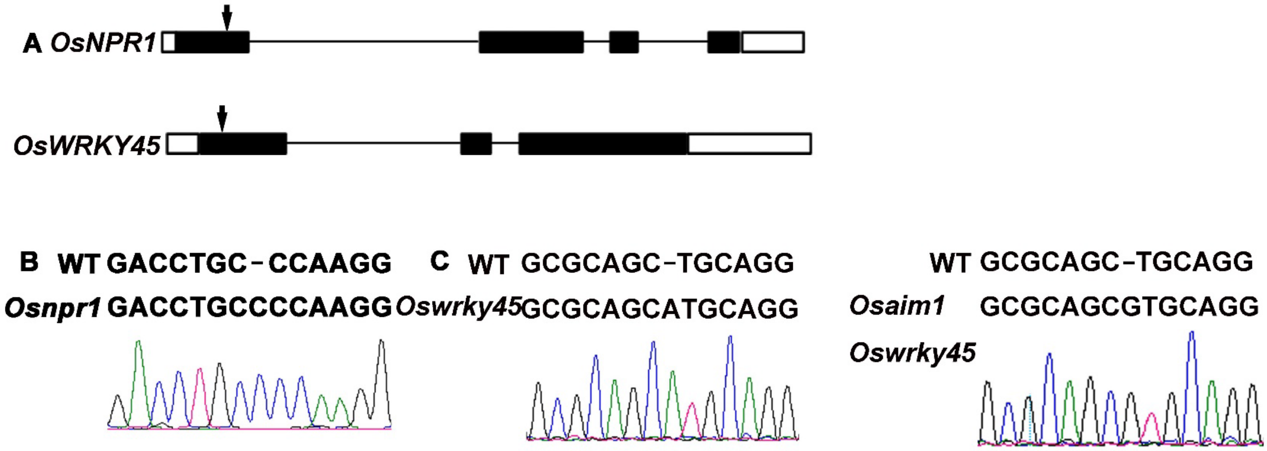

**Fig. S3 Sequence information of *NPR1* and *WRKY45* in their mutants.**

**(A)** Schematic representation of *OsNPR1* and *OsWRKY45* gene structure, and the arrows indicate the CRISPR-Cas9 target sites. **(B)** Sequence information of *OsNPR1* in *Osnpr1* mutant. **(C)** Sequence information of *OsWRKY45* in *Oswrky45* mutant and *Osaim1 Oswrky45* double mutants.

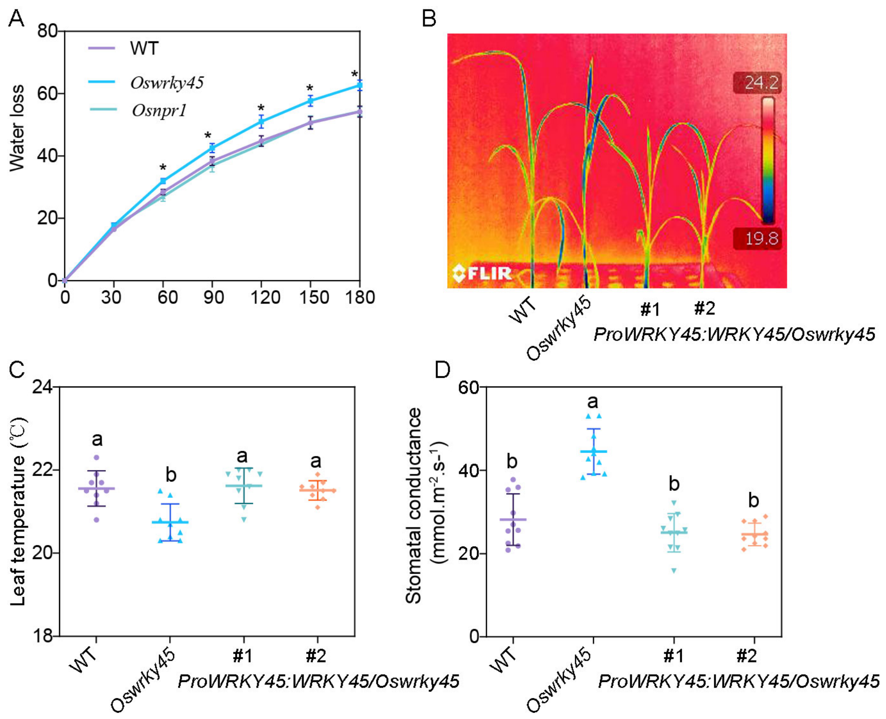

**Fig. S4 Phenotype of WT, *Oswrky45* mutant and *Oswrky45* complementation lines.**

**(A)** Water loss of detached leaves of WT, *Oswrky45* and *Osnpr1*. **(B)** Infrared images of WT, *Oswrky45* and complementation lines (*ProWRKY45:WRKY45/Oswrky45*). **(C)** Temperature of WT, *Oswrky45* and complementation lines. **(D)** Stomatal conductance of WT, *Oswrky45* and complementation lines. Error bars represent SD. Different letters show a significant difference by Tukey's test. * show a significant difference (*P<0.05, by Student's test).

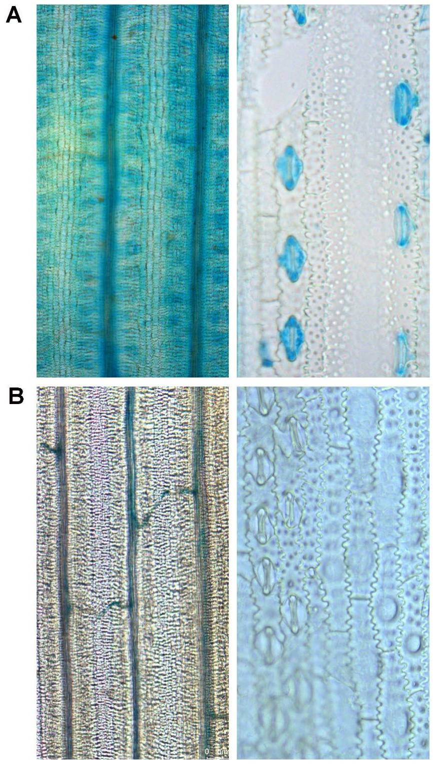

**Fig. S5** The expression of *ProWRKY45:GUS* (A) and *ProNPR1:GUS* (B) in leaf**.**

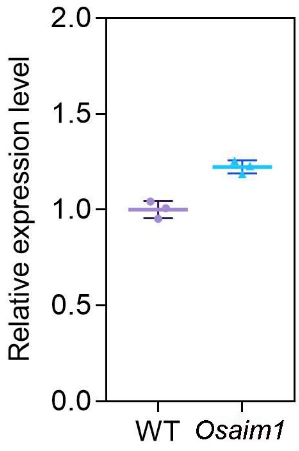

**Fig. S6** The expression level of *NPR1* in WT and *Osaim1***.** Error bars represent SD.

**Table S1.** Primer used in this study.

| Primer for qRT-PCR | Forword | Reverse |
|---|---|---|
| OsNPR1 | gcctccaccttccaggtc | tctacctcaacctt atcaaggacat |
| OsWRKY45 | aggtgcagagcgaggtca | gacgattgctgcgtcctc |
| OsDCA1 | gcgggtgttgctacagtaat | gtccttcgtcgagtcaaatagg |
| OsDST1 | gatcgacatgctcaactgga | gaaggtggtgagcgtggt |
| Primer for OsICS | | |
| RT-PCR | ccagagcaattatttcatcgga | ctccagcttcttctttatgctg |
| Mutant Verification | cctgggatcttgccgttact | cggacggtgttgatatcgc |
| Bound primer | ccacagttttcgcgatccagactg | |
| Primer for vector construction | | |
| ProWRKY45:GUS | aactgcagctgtttctcaccg gtgactca | cggggtacccctcaat ccaagcaagcaagc |
| ProNPR1:GUS | acgcgtcgacacattttt aaaaaagtgaacg | cggggtacctgcgca cccgcacccgcaccg |
| ProWRKY45:WRKY45 | cggggtaccctgtt tctcaccggtgactcat | tgctctagatcaaaa gctcaaacccataatg |
| Primer for CRISPR-Cas9 | | |
| NPR1 | ccgcgtcggcgacctgcccaagg | |
| WRKY45 | ggagctggcggcgcagctgcagg | |
